# Functional Stratification Reveals Speed-Independent Gait Impairments Beyond Chronological Age

**DOI:** 10.64898/2026.01.20.700649

**Authors:** Yuetong Wu, Xiangrui Wang, Todd M. Manini, Boyi Hu

## Abstract

**Background:** Gait is a clinically relevant indicator of functional decline in aging populations. However, most studies classify older adults by chronological rather than functional age, which may obscure early impairments detectable through kinematic profiling. This study examined whether stratifying older adults by functional status using the Short Physical Performance Battery (SPPB) enhances sensitivity in detecting gait abnormalities and instability-related compensatory patterns.

**Methods:** A total of 190 adults completed gait trials on a pressure-sensitive walkway. Twenty-eight spatial, temporal, and variability-based gait parameters were derived. Participants were categorized as young adults or older adults, who were further stratified into high- and low-functioning groups based on SPPB scores. Analysis of covariance (ANCOVA) was performed, adjusting for habitual walking speed to isolate functional effects.

**Findings:** After adjusting for speed, the low-functioning group demonstrated longer stance and double-support durations, wider step width, and greater step-to-step variability in both spatial and temporal domains compared with both the high-functioning and young reference groups. These findings indicate a compensatory, instability-driven control strategy that challenges the assumption of a “slower but steady” gait in aging. High-functioning older adults exhibited gait patterns more closely resembling those of younger adults.

**Interpretation:** Functional classification using the SPPB provided greater sensitivity than chronological age in detecting early mobility decline. Gait variability emerged as a salient biomarker of impaired neuromuscular control. Integrating quantitative gait profiling with validated functional assessments may improve early screening, targeted intervention, and fall prevention strategies.

## Introduction

The global population is aging rapidly, with life expectancy reaching 73 years in 2020 and projected to rise to 77 years by 2050 and 82 years by 2100 (Gu et al., 2021). This demographic transition challenges healthcare systems and underscores the need to understand aging’s physiological and functional impacts. With age, declines in muscle mass, strength, and central nervous system function contribute to mobility impairments and loss of independence (Ferrucci et al., 2016; Sorond et al., 2015), which in turn predict falls, hospitalization, dementia, and mortality.

Aging involves distinct changes in gait. Older adults typically walk more slowly (Masse et al., 2021; Alexander, 1996), with reduced joint range of motion and diminished neuromuscular coordination. Gait variability, including stride length, step width, and double support time, increases with age (Dingwell et al., 2017; Kang and Dingwell, 2008) due to sensory decline, slower neural processing, and compensatory strategies such as prolonged double support and wider steps (Johnson et al., 2020; Ko et al., 2010). These adaptations challenge postural control, making gait variability a sensitive marker of functional decline and an early indicator of frailty, cognitive impairment, and fall risk (Ferrucci et al., 2016).

Several studies have characterized gait in older adults, but key gaps remain. Tang et al. (2025) identified gait domains in adults aged 45–80 without assessing functional mobility. Hollman et al. reported normative data for 23 parameters for adults over 70 but excluded younger controls and did not stratify performance. Verlinden et al. (2013) grouped participants by age rather than function. Dapp et al. (2022) and Hansen et al. (2023) used the Short Physical Performance Battery (SPPB) but with limited parameters or wearable sensors. Although normative data for younger adults exist, differing protocols limit comparability.

This study builds on prior work with several methodological advances. We use SPPB (Pavasini et al., 2016) to classify participants, providing a validated functional assessment of lower-extremity performance. The SPPB captures gait differences not explained by age or sex alone (Welch et al., 2021; Lauretani et al., 2019). Our primary objective is to compare gait characteristics among older adults with varying SPPB scores and, secondarily, to benchmark these groups against younger adults to establish normative baselines. To isolate functional effects, we employed controlled protocols and a consistent walking environment (Geerse et al., 2017). Twenty-eight gait parameters spanning spatial, temporal, and variability domains were analyzed. Traditional measures such as stride length, cadence, and support times were complemented by variability metrics. We hypothesize that (1) older adults with lower SPPB scores will show greater variability and altered spatiotemporal patterns, and (2) younger adults will exhibit more stable, symmetric gait, underscoring the impact of aging on neuromuscular control. Integrating validated functional stratification with detailed gait analysis and cross-age comparison, this study clarifies mechanisms of mobility decline and informs early screening and intervention to preserve independence in aging.

## Methods Participants

This study included 259 participants, with 192 with gait data recorded of whom 190 (61 male, 129 female) were analyzed. Eligibility required completion of the SPPB with scores from 2–12, excluding those with the lowest scores (0–1) to focus on independently ambulatory adults and examine functional impairment in relation to gait (Guralnik et al., 1995). Participants maintained stable body weight (±5 lbs) for three months. The SPPB, comprising balance, an 8-ft walk, and chair stands, assessed functional performance. Older adults were classified as LOW (2–9), MOD (10–11), or HIGH (12), with higher scores indicating better function. A younger reference group (YOUNG) without functional limitations enabled age-related comparisons and establishment of normative gait values. This design facilitated detection of functional aging effects and subclinical gait changes. All procedures were approved by the University of Florida IRB, and written informed consent was obtained (IRB #87-2013).

### Gait Parameters

Gait characteristics were evaluated using a pressure-sensitive walkway (GaitRite, CIR Systems Inc., Franklin, NJ, USA), providing validated spatial and temporal gait measures. Participants walked at a self-selected pace across the walkway three times with rest periods. Parameters were extracted and averaged across sides and trials to obtain representative values. The gait parameters were categorized into spatial, temporal, and variability domains, as described in Table 2.

**Table 1.** Participant’s Basic Information Summary (N=190)

| Variable | Group | N | Mean | SD. Deviation |
| --- | --- | --- | --- | --- |
| Age (years) | LOW | 34 | 75.74 | 6.73 |
|  | MOD | 34 | 70.71 | 6.58 |
|  | HIGH | 37 | 69.11 | 6.88 |
|  | YOUNG | 85 | 42.34 | 13.16 |
|  | Total | 190 | 58.61 | 17.92 |
| Weight (kg) | LOW | 34 | 83.74 | 19.25 |
|  | MOD | 34 | 76.51 | 19.32 |
|  | HIGH | 37 | 72.08 | 14.89 |
|  | YOUNG | 85 | 75.54 | 20.10 |
|  | Total | 190 | 76.51 | 19.11 |
| Height (cm) | LOW | 34 | 167.13 | 8.16 |
|  | MOD | 34 | 165.93 | 9.83 |
|  | HIGH | 37 | 164.46 | 7.99 |
|  | YOUNG | 85 | 169.11 | 7.96 |
|  | Total | 190 | 167.28 | 8.50 |
| BMI (kg/m²) | LOW | 34 | 29.81 | 5.82 |
|  | MOD | 34 | 27.62 | 5.92 |
|  | HIGH | 37 | 26.50 | 4.16 |
|  | YOUNG | 85 | 26.33 | 6.52 |
|  | Total | 190 | 27.22 | 5.99 |

**Table 2.**
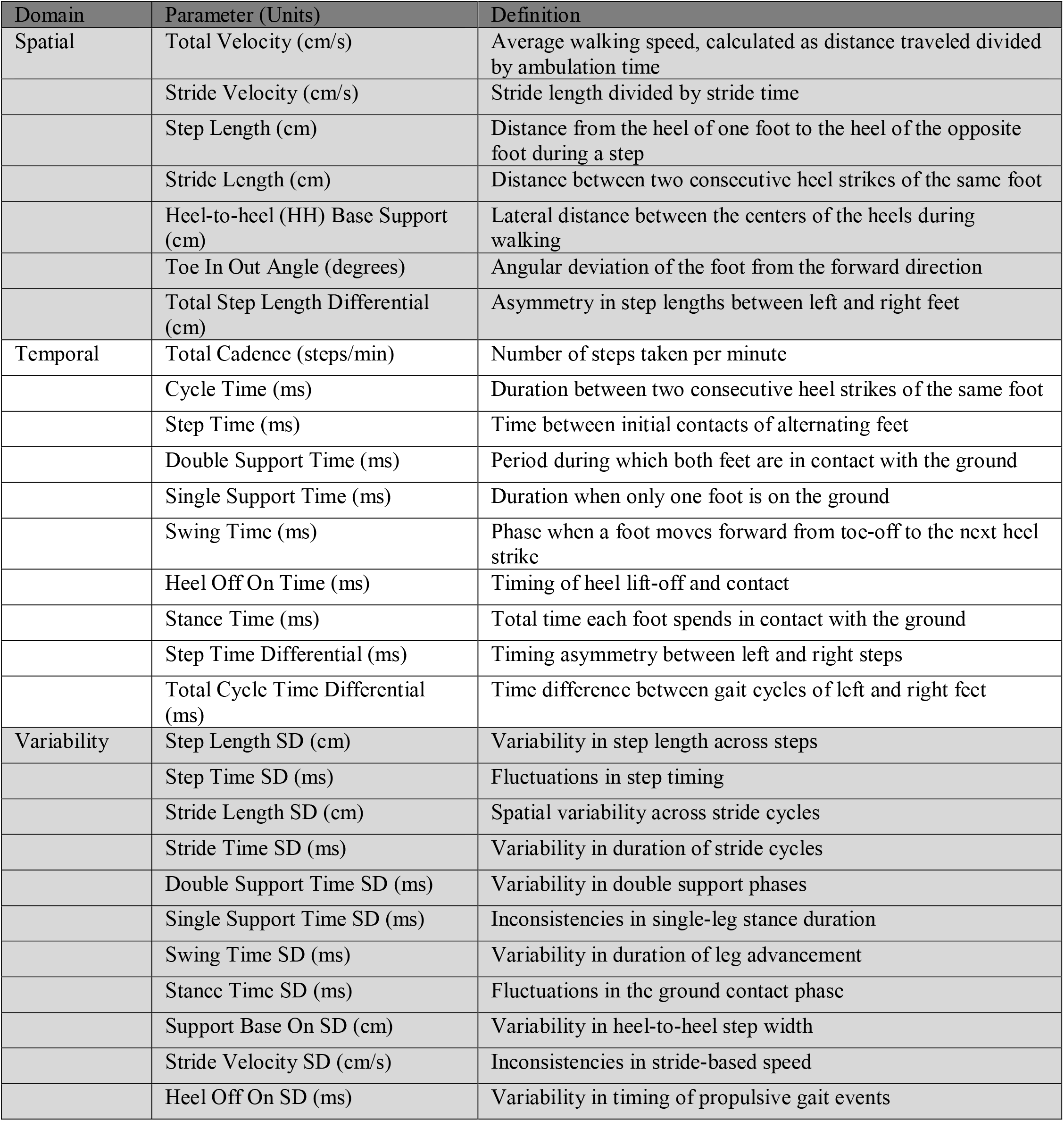
Description of Gait Parameters.

### Data Preprocessing and Statistical Analysis

Data was analyzed in SPSS Statistics 27 (IBM Corp., Armonk, NY) with significance set at p < 0.05. To account for walking speed, an analysis of covariance (ANCOVA) was performed with each participant’s mean walking velocity entered as a covariate. Adjusted group means were estimated at the overall sample mean velocity to allow comparison of groups at an equivalent walking speed. Variables showing significant main effects (p < 0.05) were further examined using Holm-adjusted post hoc tests.

## Results

### Spatial parameters

Spatial gait characteristics differed across groups after adjusting for mean walking velocity (Table 3). Total Velocity rose from 100.2 (15.9) cm/s in LOW to 127.7 (15.1) cm/s in HIGH, with YOUNG at 128.1 (20.4) cm/s. As velocity was a covariate, no inferential tests were performed. After adjustment, several spatial parameters remained significant. Step Length increased with functional ability (*F* = 4.230, *p* = 0.006): 57.4 (7.6) cm in LOW to 68.0 (7.4) cm in HIGH, and 70.0 (8.1) cm in YOUNG. HH-Base Support decreased with function (*F* = 4.945, *p* = 0.003): 11.7 (3.8) cm in LOW, 9.4 (2.8) cm in MOD, and 9.1 (2.4) cm in HIGH, with a significant LOW–YOUNG difference (+20.28%). Total Step Length Differential also differed (*F* = 3.118, *p* = 0.027): 2.8 (1.8) cm in LOW and MOD, 2.1 (1.0) cm in HIGH, and 2.0 (1.4) cm in YOUNG.

**Table 3.**
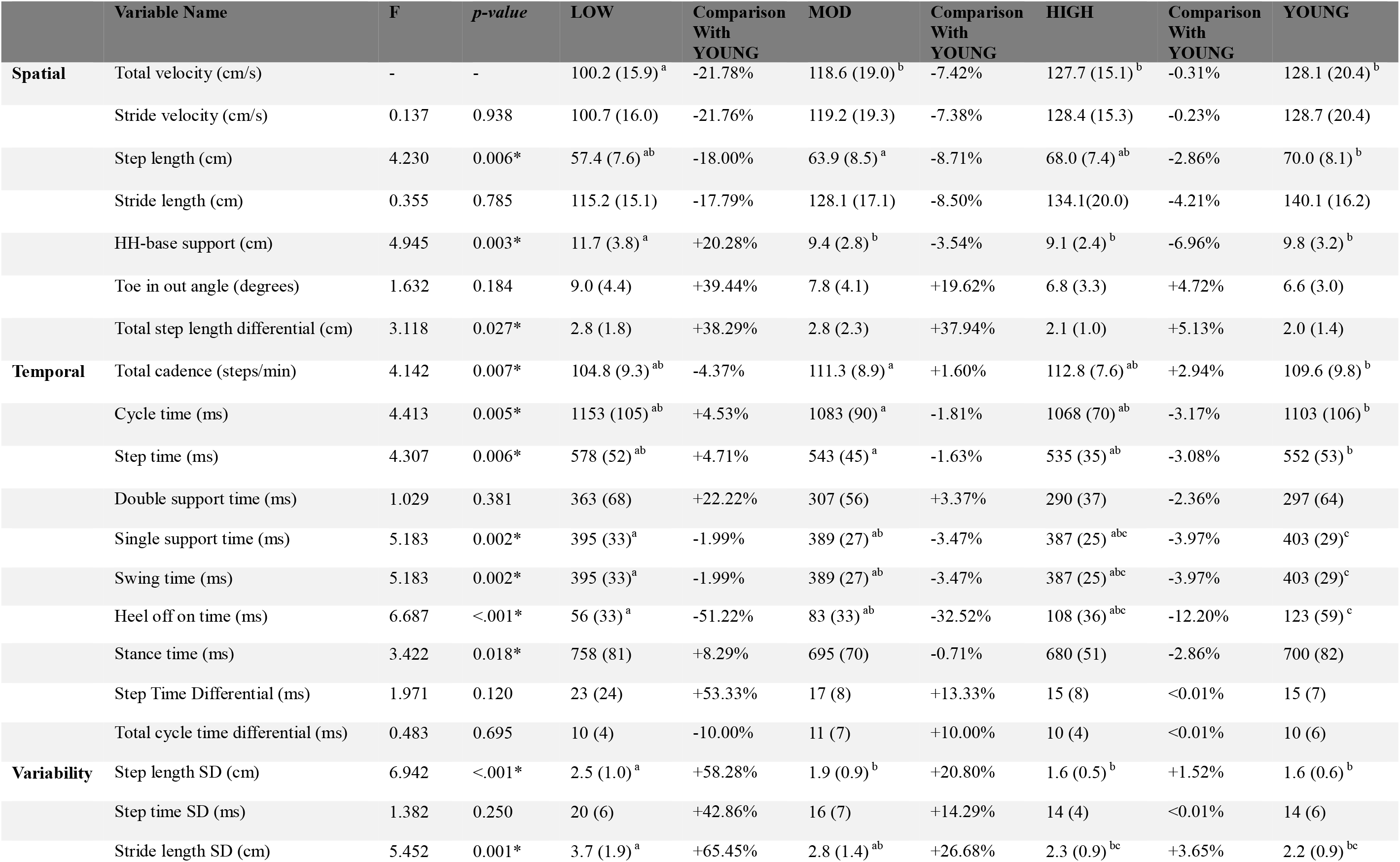
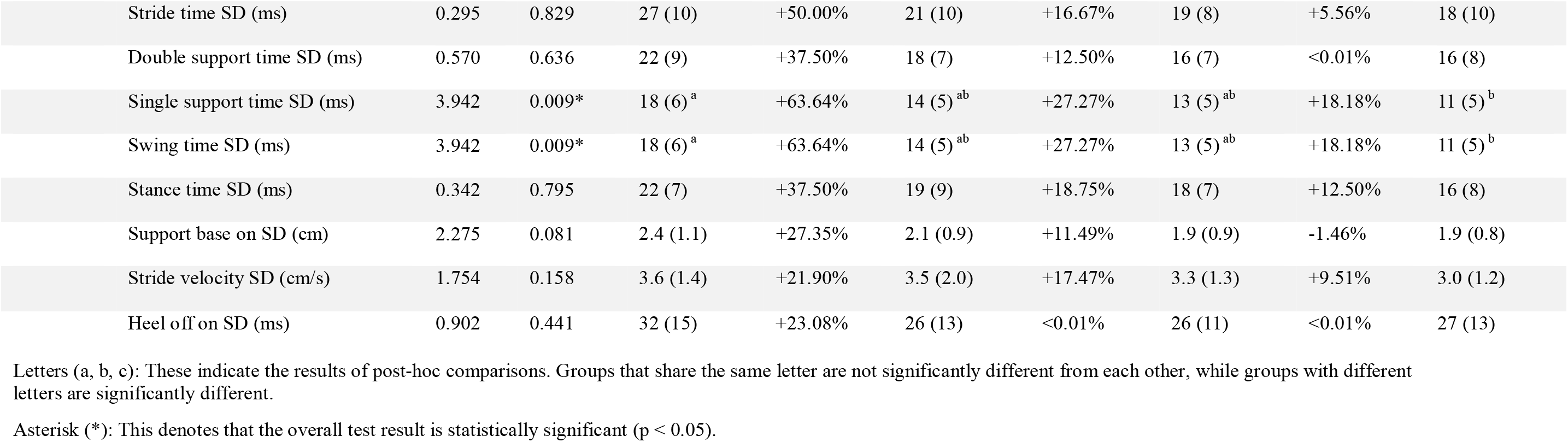
Summary of Gait Parameters between Different Groups.

|  | Variable Name | F | *p-value* | LOW | Comparison With YOUNG | MOD | Comparison With YOUNG | HIGH | Comparison With YOUNG | YOUNG |
| --- | --- | --- | --- | --- | --- | --- | --- | --- | --- | --- |
| **Spatial** | Total velocity (cm/s) | - | - | 100.2 (15.9)<sup>a</sup> | -21.78% | 118.6 (19.0)<sup>b</sup> | -7.42% | 127.7 (15.1)<sup>b</sup> | -0.31% | 128.1 (20.4)<sup>b</sup> |
|  | Stride velocity (cm/s) | 0.137 | 0.938 | 100.7 (16.0) | -21.76% | 119.2 (19.3) | -7.38% | 128.4 (15.3) | -0.23% | 128.7 (20.4) |
|  | Step length (cm) | 4.230 | 0.006* | 57.4 (7.6)<sup>ab</sup> | -18.00% | 63.9 (8.5)<sup>a</sup> | -8.71% | 68.0 (7.4)<sup>ab</sup> | -2.86% | 70.0 (8.1)<sup>b</sup> |
|  | Stride length (cm) | 0.355 | 0.785 | 115.2 (15.1) | -17.79% | 128.1 (17.1) | -8.50% | 134.1(20.0) | -4.21% | 140.1 (16.2) |
|  | HH-base support (cm) | 4.945 | 0.003* | 11.7 (3.8)<sup>a</sup> | +20.28% | 9.4 (2.8)<sup>b</sup> | -3.54% | 9.1 (2.4)<sup>b</sup> | -6.96% | 9.8 (3.2)<sup>b</sup> |
|  | Toe in out angle (degrees) | 1.632 | 0.184 | 9.0 (4.4) | +39.44% | 7.8 (4.1) | +19.62% | 6.8 (3.3) | +4.72% | 6.6 (3.0) |
|  | Total step length differential (cm) | 3.118 | 0.027* | 2.8 (1.8) | +38.29% | 2.8 (2.3) | +37.94% | 2.1 (1.0) | +5.13% | 2.0 (1.4) |
| **Temporal** | Total cadence (steps/min) | 4.142 | 0.007* | 104.8 (9.3)<sup>ab</sup> | -4.37% | 111.3 (8.9)<sup>a</sup> | +1.60% | 112.8 (7.6)<sup>ab</sup> | +2.94% | 109.6 (9.8)<sup>b</sup> |
|  | Cycle time (ms) | 4.413 | 0.005* | 1153 (105)<sup>ab</sup> | +4.53% | 1083 (90)<sup>a</sup> | -1.81% | 1068 (70)<sup>ab</sup> | -3.17% | 1103 (106)<sup>b</sup> |
|  | Step time (ms) | 4.307 | 0.006* | 578 (52)<sup>ab</sup> | +4.71% | 543 (45)<sup>a</sup> | -1.63% | 535 (35)<sup>ab</sup> | -3.08% | 552 (53)<sup>b</sup> |
|  | Double support time (ms) | 1.029 | 0.381 | 363 (68) | +22.22% | 307 (56) | +3.37% | 290 (37) | -2.36% | 297 (64) |
|  | Single support time (ms) | 5.183 | 0.002* | 395 (33)<sup>a</sup> | -1.99% | 389 (27)<sup>ab</sup> | -3.47% | 387 (25)<sup>abc</sup> | -3.97% | 403 (29)<sup>c</sup> |
|  | Swing time (ms) | 5.183 | 0.002* | 395 (33)<sup>a</sup> | -1.99% | 389 (27)<sup>ab</sup> | -3.47% | 387 (25)<sup>abc</sup> | -3.97% | 403 (29)<sup>c</sup> |
|  | Heel off on time (ms) | 6.687 | <.001* | 56 (33)<sup>a</sup> | -51.22% | 83 (33)<sup>ab</sup> | -32.52% | 108 (36)<sup>abc</sup> | -12.20% | 123 (59)<sup>c</sup> |
|  | Stance time (ms) | 3.422 | 0.018* | 758 (81) | +8.29% | 695 (70) | -0.71% | 680 (51) | -2.86% | 700 (82) |
|  | Step Time Differential (ms) | 1.971 | 0.120 | 23 (24) | +53.33% | 17 (8) | +13.33% | 15 (8) | <0.01% | 15 (7) |
|  | Total cycle time differential (ms) | 0.483 | 0.695 | 10 (4) | -10.00% | 11 (7) | +10.00% | 10 (4) | <0.01% | 10 (6) |
| **Variability** | Step length SD (cm) | 6.942 | <.001* | 2.5 (1.0)<sup>a</sup> | +58.28% | 1.9 (0.9)<sup>b</sup> | +20.80% | 1.6 (0.5)<sup>b</sup> | +1.52% | 1.6 (0.6)<sup>b</sup> |
|  | Step time SD (ms) | 1.382 | 0.250 | 20 (6) | +42.86% | 16 (7) | +14.29% | 14 (4) | <0.01% | 14 (6) |
|  | Stride length SD (cm) | 5.452 | 0.001* | 3.7 (1.9)<sup>a</sup> | +65.45% | 2.8 (1.4)<sup>ab</sup> | +26.68% | 2.3 (0.9)<sup>bc</sup> | +3.65% | 2.2 (0.9)<sup>bc</sup> |

|  |  |  |  |  |  |  |  |  |  |
| --- | --- | --- | --- | --- | --- | --- | --- | --- | --- |
| Stride time SD (ms) | 0.295 | 0.829 | 27 (10) | +50.00% | 21 (10) | +16.67% | 19 (8) | +5.56% | 18 (10) |
| Double support time SD (ms) | 0.570 | 0.636 | 22 (9) | +37.50% | 18 (7) | +12.50% | 16 (7) | <0.01% | 16 (8) |
| Single support time SD (ms) | 3.942 | 0.009* | 18 (6)<sup>a</sup> | +63.64% | 14 (5)<sup>ab</sup> | +27.27% | 13 (5)<sup>ab</sup> | +18.18% | 11 (5)<sup>b</sup> |
| Swing time SD (ms) | 3.942 | 0.009* | 18 (6)<sup>a</sup> | +63.64% | 14 (5)<sup>ab</sup> | +27.27% | 13 (5)<sup>ab</sup> | +18.18% | 11 (5)<sup>b</sup> |
| Stance time SD (ms) | 0.342 | 0.795 | 22 (7) | +37.50% | 19 (9) | +18.75% | 18 (7) | +12.50% | 16 (8) |
| Support base on SD (cm) | 2.275 | 0.081 | 2.4 (1.1) | +27.35% | 2.1 (0.9) | +11.49% | 1.9 (0.9) | -1.46% | 1.9 (0.8) |
| Stride velocity SD (cm/s) | 1.754 | 0.158 | 3.6 (1.4) | +21.90% | 3.5 (2.0) | +17.47% | 3.3 (1.3) | +9.51% | 3.0 (1.2) |
| Heel off on SD (ms) | 0.902 | 0.441 | 32 (15) | +23.08% | 26 (13) | <0.01% | 26 (11) | <0.01% | 27 (13) |

Letters (a, b, c): These indicate the results of post-hoc comparisons. Groups that share the same letter are not significantly different from each other, while groups with different letters are significantly different.
Asterisk (\*): This denotes that the overall test result is statistically significant ( $p < 0.05$ ).

### Temporal parameters

Temporal gait features also varied by function (Table 3). Total Cadence differed significantly (*F* = 4.142, *p* = 0.007), ranging from 104.8 (9.3) steps/min in LOW to 112.8 (7.6) in HIGH, with MOD showing pairwise effects and YOUNG at 109.6 (9.8) steps/min. Cycle Time and Step Time followed similar patterns (*F* = 4.413, *p* = 0.005; *F* = 4.307, *p* = 0.006), with MOD significantly lower than YOUNG. Single Support and Swing Time differed (*F* = 5.183, *p* = 0.002), shorter in LOW and longer in YOUNG. Heel Off On Time increased with function (*F* = 6.687, *p* < 0.0011), from 56 (33) ms in LOW to 108 (36) in HIGH, and 123 (59) in YOUNG. Stance Time also differed (*F* = 3.422, *p* = 0.018), from 758 (81) ms in LOW to 680 (51) in HIGH.

### Variability parameters

Gait variability showed significant group differences (Table 3). Step Length SD differed (*F* = 6.942, *p* < 0.001), highest in LOW (2.5 (1.0) cm), followed by MOD (1.9 (0.9)), HIGH (1.6 (0.5)), and YOUNG (1.6 (0.6)). Only LOW differed significantly from YOUNG (+58.28%). Stride Length SD also varied (*F* = 5.452, *p* = 0.001), with LOW showing a 65.45% increase versus YOUNG. Single Support Time SD and Swing Time SD differed (*F* = 3.942, *p* = 0.009), decreasing from 18 (6) ms in LOW to 13 (5) ms in HIGH, with YOUNG at 11 (5). Other variability measures were non-significant (Table 3, Figure 1).

**Figure 1.**
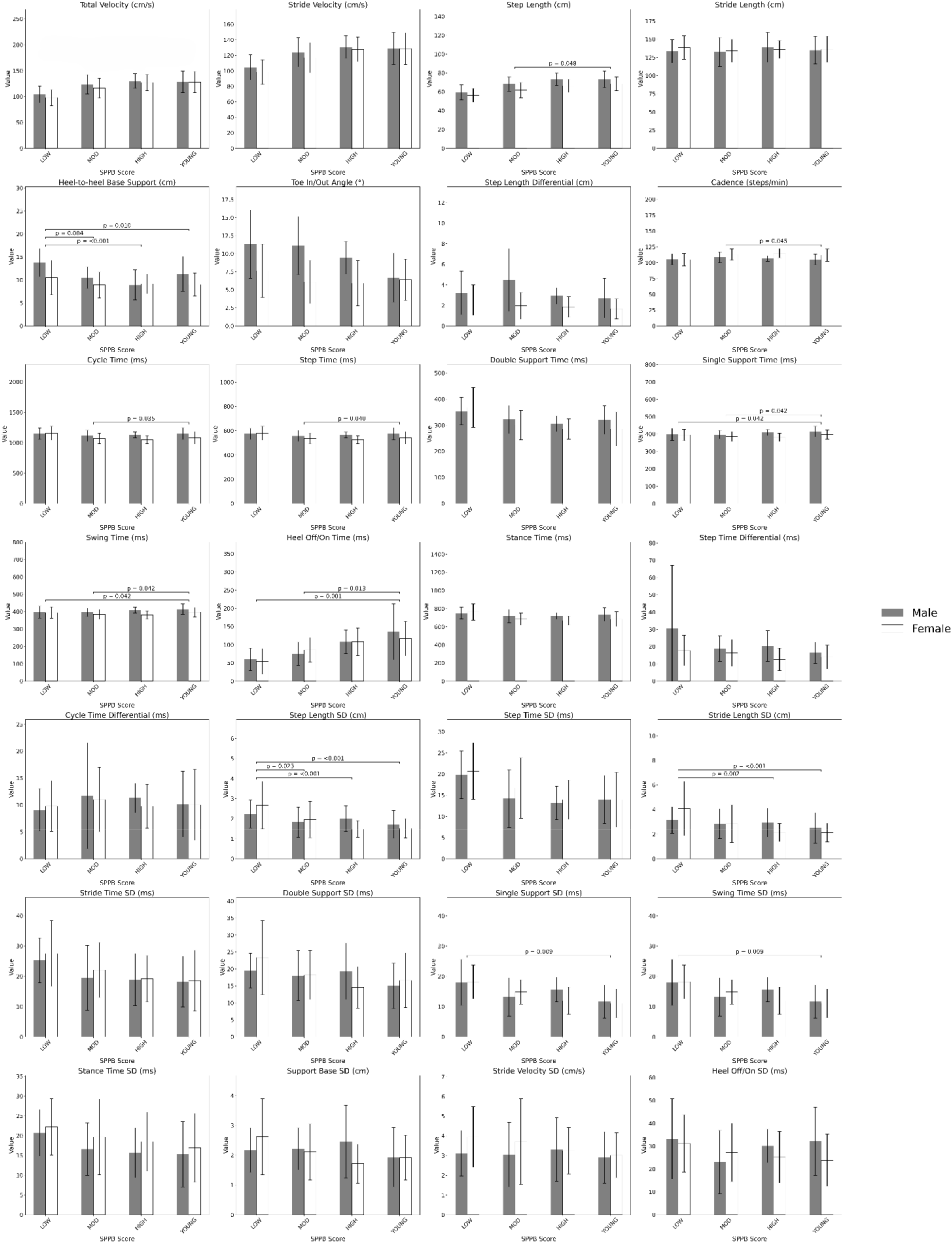
Comparison of Gait Parameters Across SPPB Categories

## Discussion

The primary objective of this study was to analyze gait characteristics across adults with different functional levels. In the context of health prevention and early intervention, individual differences in function are often more clinically meaningful than age alone. Although prior research (Dapp et al., 2022), used the SPPB to stratify older adults, our study extends this approach by incorporating a richer set of gait parameters for a more comprehensive, high-resolution profile. By including a younger reference group and a wide range of spatial, temporal, and variability measures under the same protocol, this study provides a broader quantitative perspective on age- and function-related gait adaptations and highlights opportunities for tailored, function-based mobility interventions. Given established links between gait, falls, disability, and mortality, our detailed gait profiles may help identify subtle impairments before overt clinical decline.

### Spatial and Temporal Parameters

Spatial parameters showed robust functional stratification across groups. Total (gait) Velocity, often described as the “sixth vital sign” in geriatrics (Middleton et al., 2015), was used as a covariate to control for walking speed and was not included in inferential testing. Descriptively, velocity increased with higher functional ability, consistent with evidence linking gait speed to global health and physiological reserve. Prior work indicates that a usual gait speed below 1 m/s identifies individuals at elevated risk of morbidity and mortality (Studenski et al., 2011; Cesari et al., 2005).

Although cadence is typically classified as a temporal and step length as a spatial parameter, both jointly determine walking speed. In this study, both varied significantly with functional ability, revealing compensatory gait strategies across performance levels. Total Cadence rose from 104 ± 9.3 steps/min in LOW to 112.8 ± 7.6 steps/min in HIGH, while YOUNG showed slightly lower cadence (109.6 ± 9.8 steps/min), likely reflecting longer steps. These results are consistent with Hollman et al. (2011), who observed shorter step lengths and altered cadence with advancing age. However, our findings extend this work by demonstrating that such spatial adaptations persist even after controlling for mean walking velocity. Thus, the observed reductions in step length cannot be attributed merely to slower pace but rather represent structural adaptations in movement control. This interpretation supports the view that shorter steps in lower-functioning adults are not voluntary pacing adjustments but reflect constrained motor output, marking an early biomechanical indicator of reduced neuromuscular efficiency.

HH-Base Support (step width), which differed significantly across groups (*p* =.003), also reflected compensatory stability strategies. Step width increased from 9.1 ± 2.4 cm (HIGH) to 9.4 ± 2.8 cm (MOD) and 11.7 ± 3.8 cm (LOW), compared to 9.8 ± 3.2 cm in YOUNG. The 20.28% wider base in LOW remained after adjusting for speed, challenging the idea that base widening results solely from slower gait. Instead, it suggests structural reorganization to maintain stability before overt instability appears. Although widening the base aids balance, it raises energy cost and alters mechanics. Prior studies have described this as a compensatory strategy: Menz et al. (2003) observed that older adults tend to widen their base of support to preserve balance in response to age-related sensory and motor decline, and Bauby and Kuo (2000) demonstrated that a wider base improves stability but demands greater mechanical work and energy expenditure during walking. Building on this evidence, our findings reposition base widening as a fundamental marker of degraded postural control rather than a secondary adaptive response.

Step Length Differential, quantifying interlimb asymmetry, further illuminated functional gait control. While Zadik et al. (2022) found no age-related asymmetry, our function-based stratification showed that interlimb asymmetry, as measured by step length differential (*p* =.03), increased with lower physical function among older adults, from 2.8 ± 1.8 cm in the LOW group to 2.1 ± 1.0 cm in the HIGH group. Notably, the LOW group exhibited a significant increase of +38.29% compared to the baseline. This discrepancy highlights a key limitation in age-based grouping: it may obscure functionally relevant differences in gait. By examining asymmetry through a functional lens, our findings indicate that gait asymmetry does indeed increase in individuals with lower function. These results suggest that asymmetry may emerge not merely with age, but because of declining physical capacity, reinforcing its potential utility as a sensitive marker of early functional deterioration.

Taken together, the spatial block shows that even the expected age-related adaptations, shorter steps, wider base of support, and greater asymmetry, persist under speed normalization, indicating that these are not trivial speed-driven artifacts but structural signatures of degraded control that precede and likely contribute to the stronger instability patterns observed in the temporal and variability domains.

Temporal components such as Cycle Time and Step Time decreased with increasing function across groups (*p* = 0.005 and *p* = 0.006). Interestingly, the YOUNG group did not exhibit the shortest durations as might be expected. Shorter cycle times are generally associated with greater gait efficiency and reduced fall risk (Verghese et al., 2009). Instead, baseline values fell between those of the MOD and LOW groups, likely reflecting our use of functional rather than age-based stratification. The HIGH group, comprising older adults with better SPPB scores (Verlinden et al., 2013), showed stride lengths comparable to YOUNG, reinforcing that functional ability more strongly shapes gait patterns. Similarly, Hollman et al. (2011) reported minimal step-time differences across age groups, suggesting it may be an insensitive marker of gait decline and balance impairment.

Single Support Time and Swing Time increased slightly with function (*p* =.020), from 395 ± 33 ms in the LOW group to 387 ± 25 ms in the HIGH group. These findings align with those of Sung (2018), who reported that older adults with lower bone mineral density exhibited increased double support times during walking, suggesting a compensatory mechanism for diminished postural control. While Sung’s study focused on limb dominance and bone health, our function-based stratification reveals that such compensatory strategies are broadly evident in lower-functioning individuals. The inverse pattern of single and double support durations across functional groups underscores the interdependence of these phases and highlights their value as sensitive markers of gait adaptation in aging populations.

Heel Off On Time, which reflects the terminal stance to pre-swing phase, increased significantly with function (*p <*.*001*), starting from 123 ± 53 ms in the baseline. The HIGH group showed a value of 108 ± 36 ms, followed by the MOD group, and decreasing to 56 ± 33 ms in the LOW group. The significant difference between the LOW and MOD group with the baseline reached a -51.22% and –32.52% decrease, respectively. This parameter may reflect improved push-off mechanics and ankle control, where longer heel-off times indicate more complete utilization of the trailing limb during propulsion. Efficient push-off contributes to forward momentum and gait smoothness, which are often compromised in lower-functioning individuals with plantar flexor weakness or joint stiffness (Judge et al., 1996).

In contrast, Stance Time (*p=0*.*18*), which captures the duration of foot contact during the gait cycle did not show a consistent decreasing trend. For older adults, while the value for the MOD and HIGH groups was similar, the LOW groups exhibited the highest stance time of 0.758 seconds. Longer stance times in this group likely reflected cautious walking behavior and impaired neuromuscular control, where there is increased ground contact time?

### Variability Parameters

Gait variability, defined as stride-to-stride fluctuations in spatiotemporal features, offers key insight into neuromuscular control and dynamic stability. Its importance has been repeatedly demonstrated in literature. Hollman et al. (2011) and Brach et al. (2001) showed that increased gait variability is strongly associated with fall risk, cognitive impairment, and loss of neuromotor automaticity. Specifically, Hollman et al. (2011) reported that higher stride-to-stride fluctuations predict functional decline and fall propensity in aging adults. While gait speed is a common clinical marker, our variability analysis identifies subclinical motor-control deficits that may precede overt mobility disability. Linking specific variability measures to SPPB domains, strength, balance, and coordination provides mechanistic insight into affected components of lower-extremity function and informs domain-targeted interventions.

In our analysis, variability metrics showed significant stratification by functional level. Step Length SD decreased (*p* <.001) from 2.5 ± 1.0cm in the LOW group to 1.6 ± 0.5 cm in the HIGH group, with the lowest value (1.6 ± 0.6 cm) in the YOUNG group. In addition, Stride Length SD also showed a strong functional decrease in our study (*p* <.001), from the LOW to HIGH and again lowest in YOUNG groups. This aligns well with findings by Dapp et al. (2022) who reported a significant increase in stride length variation across robust, transient, and frail older adults, with values rising from 3.2% in the robust group to 5.4% in the frail group using the SPPB classification. These findings are consistent with the premise that heightened spatial variability reflects impaired sensorimotor integration and increased fall risk (Hausdorff et al., 2001; Brach et al., 2007).

Notably, while Larsson et al. (2016) found no significant differences in swing time SD, our study revealed a clear stratification (*p* =.009), with Swing Time SD decreasing across groups. Specifically, the LOW group (18 ± 6 ms) exhibited greater variability than the baseline (11 ± 5 ms), corresponding to 63.64% and 27.27% increases, respectively. Although This discrepancy may be due to the fact that our study grouped participants based on physical function, whereas Larsson et al. (2016) grouped participants by vestibular status. Swing time SD may be more sensitive to overall physical function and neuromotor control than to vestibular problems alone, making it a broader and more useful indicator of gait instability in older adults living in the community. Similarly, Single Support Time SD showed identical results, suggesting that greater balance confidence and control allows for more consistent unilateral stance.

Taken together, these findings position gait variability as a powerful marker of functional status, often more sensitive than mean-level gait features. Compared to studies assessing variability in isolation or clinical cohorts, our work provides a comprehensive variability profile across the healthy aging spectrum. It also highlights the superiority of functional over chronological classification, as variability measures consistently mirror gradations in physical capability. These metrics should be incorporated into mobility screening and fall risk assessments to detect early decline in older adults who may otherwise appear high functioning.

## Limitations

While this study provides a comprehensive analysis of gait characteristics across functional groups, several limitations exist. First, although the SPPB offers a validated measure of lower-extremity function, the design limits inference about longitudinal change or clinical prediction. We did not assess whether specific gait parameters predict falls, hospitalization, or mortality; future longitudinal work should address this. Second, gait data were collected under controlled laboratory conditions using a pressure-sensitive walkway. Although this ensured standardization, it may not fully represent real-world gait. Ecologically valid assessments using wearable sensors could enhance generalizability, as individuals often adopt more cautious strategies in daily environments. Thus, observed performance may overestimate stability, particularly in lower-functioning groups. Future studies should incorporate free-living assessments to better reflect everyday mobility demands.

## Conclusion

This study advances understanding of gait adaptations in aging by comparing gait characteristics among older adults stratified by SPPB scores and benchmarking against a younger reference group. Using 28 gait parameters, we identified performance-related differences in both mean and variability measures, with lower-functioning adults showing slower, more asymmetric gait and greater variability, suggesting early neuromuscular decline. The younger group provided normative baselines that clarified functional aging patterns. Overall, findings emphasize the distinction between chronological and functional age and support incorporating function-based gait assessments into geriatric screening and prevention strategies.

## Data availability

Data will be made available upon reasonable request.

## Acknowledgments Funding

The Metabolic Costs of Daily Activity in Older Adults Study is funded by the National Institutes of Health (NIH)/National Institute on Aging (NIA) (R01AG042525). The research is also partially supported by the Claude D. Pepper Older Americans Independence Centers at the University of Florida (1P30AG028740)

